# GPCR Evolution Database

**DOI:** 10.64898/2026.09.08.750181

**Authors:** Berkay Selçuk, Ogün Adebali

## Abstract

GPCRs make up the largest family of human membrane proteins and of drug targets. Decades of experimental and structural work have revealed how these receptors operate at the molecular level, but this work has covered only a fraction of the superfamily, leaving the majority of receptors underexplored. To address this problem, we developed the GPCR Evolution Database, gpcrevolution.org, an open-access resource that makes high-quality evolutionary analysis of GPCRs available to every laboratory. Through our database, each human receptor can be examined against its own orthologs, where per-residue conservation reveals the sites that evolution has protected throughout that receptor’s history. These orthologous lineages can then be compared with their paralogs within the same GPCR family, to distinguish residues shared across paralogous lineages, which likely support ancestral functions, from the lineage-specific residues that underlie functional differentiation between receptor subtypes. The database provides ortholog sets, multiple sequence alignments, phylogenetic trees and per-residue conservation scores for over 800 human GPCRs. The resource presents these through interactive conservation plots, sequence logos and snake plots, and provides tools for comparing conservation across two or more paralogous lineages. Overall, the GPCR Evolution Database provides the data and tools for researchers to examine GPCR function through an evolutionary lens, allowing molecular insights from well-studied receptors to be extended to their underexplored relatives.

## Introduction

G protein-coupled receptors (GPCRs) constitute the largest protein superfamily in the human genome^1^. They sense a wide range of external and internal stimuli and induce necessary downstream pathways that regulate numerous physiological processes. Their central role in cellular signaling makes GPCRs prime drug targets^2^, as therapeutics are often designed to artificially modulate GPCR signaling to achieve specific physiological outcomes. Over the last several decades, experimental and structural approaches have been highly successful in revealing how GPCRs operate at the molecular level. However, these efforts have focused on a limited number of receptors or receptor subfamilies, leaving many GPCRs uncharacterized.

While studying novel GPCR subtypes experimentally can be challenging, evolutionary analyses could be used to bridge the gap between well-studied and understudied receptors. Evolution, as the longest ongoing experiment, continually tests GPCR sequences through mutations and eliminates variants that reduce organismal fitness. For a given GPCR subtype (e.g. orthologs of beta-2 adrenoreceptor), sites that are resistant to mutagenesis often play significant role in receptor functioning and conserved within orthologs. Furthermore, comparisons across paralogous lineages (e.g. comparing orthologs of beta-1 and beta-2 adrenoreceptors) then highlight common residues that support shared ancestral functions, as well as lineage-specific residues that drive functional diversification among GPCR subtypes^3^. Thus, by using comparative evolutionary analyses, we can place experimental findings into an evolutionary context, allowing molecular insights from well-studied receptors to be generalized across GPCR subfamilies and families by tracing how functionally associated residues are conserved, or diversified throughout evolutionary history. In addition, evolutionary analyses can independently reveal molecular hotspots that form the molecular basis of shared and lineage-specific GPCR functions^4,5^, providing a complementary framework for generating experimentally testable hypotheses. However, conducting high-quality evolutionary analyses typically requires substantial expertise in analyzing large amounts of sequencing data and access to computational resources.

Currently, GPCRdb^6^ is the primary resource that enables comparative analysis of GPCR sequences, and it provides an invaluable compilation of curated information on human GPCRs, putative orthologs, residue annotations, receptor structures, physiological ligands, drugs, and mutational data. However, despite its broad utility, GPCRdb is not primarily designed for performing evolutionary analyses. First, the orthologous sets are imported directly from UniProtKB^7^ rather than inferred through a dedicated phylogenetic pipeline, and they can lack the evolutionary depth needed for robust comparative analyses. Second, the multiple sequence alignments are generated by extending structure-based reference alignments from human receptors, rather than by explicitly optimizing homology across diverse taxa, which can reduce their suitability for high-resolution evolutionary analyses. Third, phylogenetic relationships among orthologs are not readily available to the user and could only be inferred by using neighbor-joining trees instead of maximum-likelihood or Bayesian approaches. Finally, GPCRdb does not provide tools for systematic identification of residue-level conservation patterns across paralogous lineages to distinguish shared ancestral features from lineage-specific adaptations.

Here, we present the GPCR Evolution Database (gpcrevolution.org), an open-access resource that addresses this gap by providing evolutionary annotations and interactive analysis tools for over 800 human GPCRs and their orthologs. Building on a curated collection of receptor-specific ortholog sets, multiple sequence alignments, and phylogenetic trees from our companion evolutionary study^8^, the database quantifies per-residue conservation within each orthologous lineage and maps these scores onto bar plots, sequence logos, and two-dimensional snake plots. Beyond per-receptor views, our platform offers tools to compare conservation patterns across multiple paralogous receptor lineages within the same GPCR family. Together, these features make comparative evolutionary analysis of GPCRs accessible without specialized phylogenetic expertise or dedicated computational resources, allowing researchers to interpret molecular findings in an evolutionary context and generate experimentally testable hypotheses.

## Results

### Homepage Organization

The homepage of the GPCR Evolution Database serves as the main access point to all resources. The central search bar allows users to search for their receptor of interest by simply typing gene or receptor name to navigate to its dedicated receptor page, which contains information about its orthologs, phylogenetic tree, alignments, and residue conservation. A secondary search bar with the identical capability is also present in the site header, making receptor lookup available from any page. Below the search bar, we present the key database statistics such as the number of human GPCRs covered, the total number of ortholog sequences, and the number of integrated analysis tools, which will be updated as the database gets updated.

The global navigation menu in the header provides direct access to a “Tools” dropdown, which links to three analysis tools for comprehensive evolutionary comparisons across human GPCRs. In addition to these tools, the menu includes links to “Contact,” “FAQ,” and “Cite Us.” The Contact page provides information for feedback and collaboration inquiries, the FAQ page addresses common questions and explains the underlying data and how to interpret the outputs, and the Cite Us page lists the publications associated with the database along with the recommended citations and references for users who wish to acknowledge the resource in their work.

### Receptor Page

For each human receptor in the database, the receptor page first summarizes key metadata, including its GPCR class/family, the number of orthologs identified across species, the inferred last common ancestor, which indicates how far back the receptor’s orthologous lineage can be traced, and the corresponding UniProt^7^/GPCRdb^6^ identifier. This receptor specific page enables researchers to integrate evolutionary information into their work for their receptor of interest without needing to perform sequence searches, generate alignments and gene trees. For a given human receptor, we quantified per-residue conservation across orthologs, and this information is visualized in three different ways (Fig. 1a-c). First, at the top, conservation percentages within orthologs are presented as a bar plot displaying percentages (Fig. 1a). This is followed by the sequence logo visualization highlighting the amino-acid distribution at each site (Fig. 1b). Lastly, we utilized snakeplot visualization (imported from GPCRdb^6^) to map conservation percentages onto the two-dimensional receptor topology (Fig. 1c). Together, these views allow users to link residue conservation to structural regions and functional motifs. In a separate panel, the page also displays the phylogenetic tree of orthologous sequences alongside the multiple-sequence alignment utilized for calculation of residue conservation (Fig. 1d), enabling users to see how specific variants are distributed across clades and where substitutions arise along the gene tree. All visualizations are accompanied by download buttons, enabling users to export plots in SVG format for further customization and download the underlying data associated with that plot.

**Figure 1.**
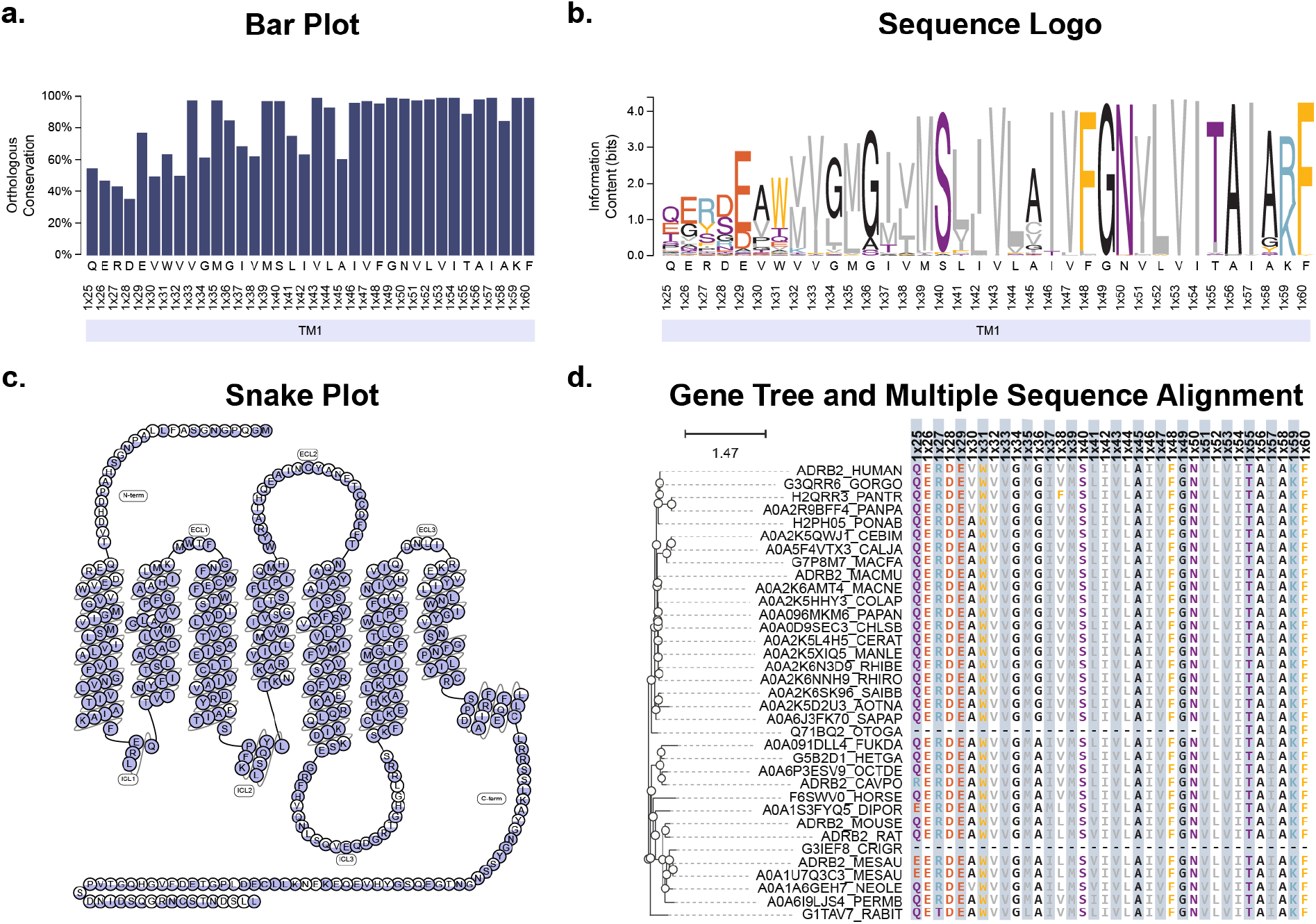
Contents of the receptor page for the beta-2 adrenergic receptor (ADRB2). a. Per-residue conservation percentages within orthologs presented as a bar plot. b. The same conservation information shown as a sequence logo, highlighting the amino acid distribution at each site. c. Snake plot representation, in which conservation percentages are mapped onto the two-dimensional receptor topology. d. Combined panel showing the gene tree of orthologous sequences alongside the multiple sequence alignment columns corresponding to human receptor residues. This page can be reached at https://gpcrevolution.org/receptor?gene=ADRB2.

### Tools

#### Differential Residue Conservation

This tool compares residue conservation patterns between two human GPCRs and their orthologs. Users select two receptors from the same GPCR family by gene name and choose a conservation threshold (default 90%), which defines which positions are considered conserved within each receptor’s ortholog set. For every aligned position, the tool combines ortholog-based conservation scores with structural region and GPCRdb generic numbering, and categorizes the site as a common residue (conserved in both receptors with similar amino-acid type, BLOSUM80 score greater than 1), specifically conserved in one receptor, or specifically conserved in both but with different preferred amino acids. The primary output is an interactive, sortable table that, for each mapped position in both receptors, reports the human residue number, human amino acid, conserved amino acid, conservation percentage, structural region and GPCRdb number, together with its conservation categories.

In addition to the table, the tool provides two additional visualizations that use the same residue categories. First, a dual sequence-logo view displays aligned sequence logos for side-by-side comparison, highlighting the sequence variability within each orthologous lineage further. Second, we utilized snake plots (imported from GPCRdb) to overlay the conservation categories onto the two-dimensional receptor topology, with interactive annotations and exportable SVGs. Together, these outputs enable users to explore common residues that are likely involved in shared ancestral functions, and receptor specific residues that are likely to create the molecular basis for functional differentiation between two paralogous lineages, moving beyond comparison of paralogous human GPCR sequences.

It is noteworthy that, in this specific example we provided in Figure 2, our tool was able to replicate our previous analysis^5^ and identify differentially conserved residues within the dimerization interface of Beta-2 adrenoceptor. We demonstrated the co-evolution between V1x33 and S1x40, and their involvement of constitutive internalization.

**Figure 2.**
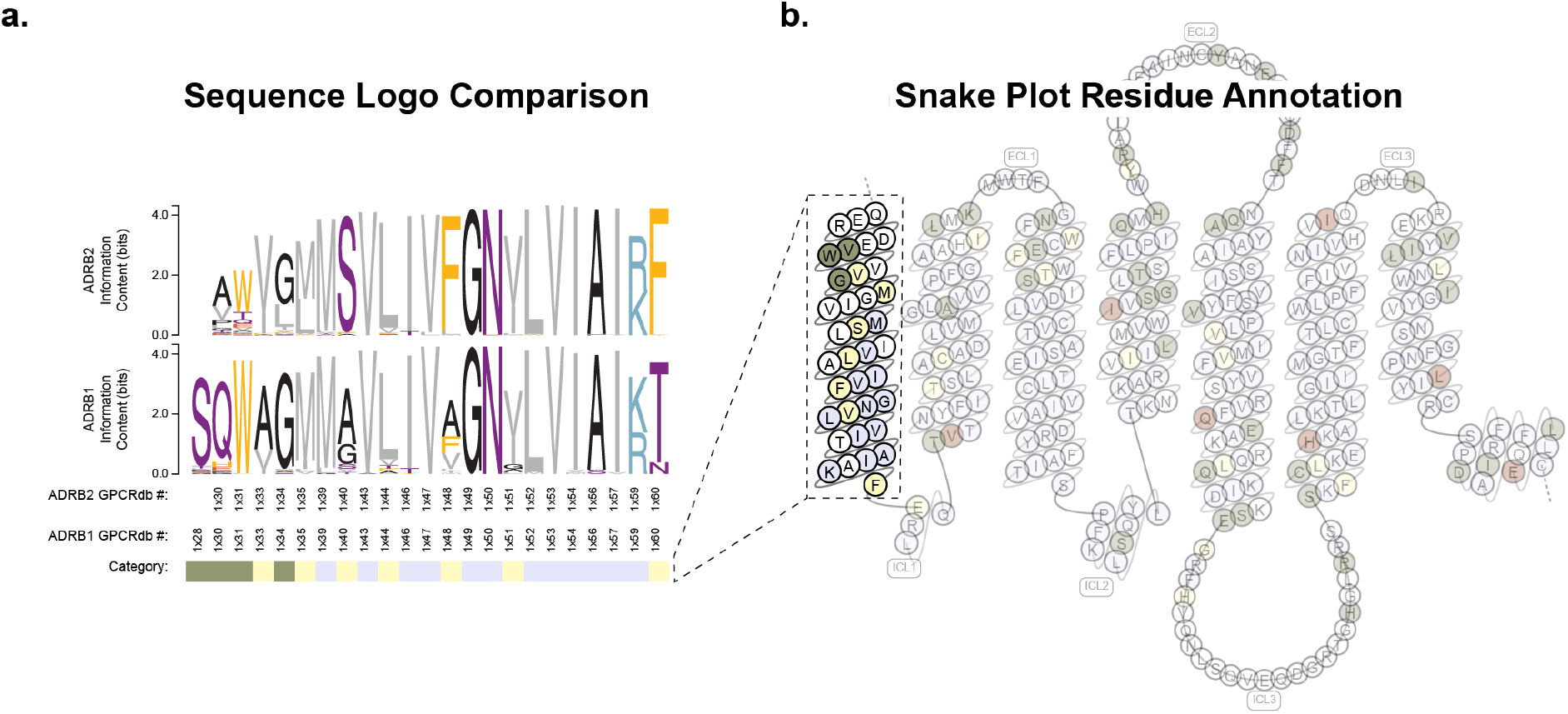
Differential conservation analysis reveals specifically conserved residues. a. Sequence logo comparison of residue conservation patterns. Beta-1 and Beta-2 adrenoreceptor residue conservation within their orthologs were compared in logo representation for the residues at transmembrane helix 1 (TM1). Below logos GPCRdb numbering of human receptors and residue categories are shown. Blue indicates common residues that are shared between two lineages, yellow indicates specific conservation for Beta-2 adrenoreceptor (ADRB2) and green indicates specific conservation for Beta-1 adrenoreceptor (ADRB1). b. The same evolutionary information was shown in snakeplot representation

#### Multi-Receptor Comparison

The multi-receptor comparison tool extends the differential conservation analysis to more than two receptors within the same GPCR class. Users select a single reference receptor and one or more target receptors (all from the same GPCR family, e.g. Class A) and can optionally restrict the analysis to specific residue numbers of the reference receptor. The tool then aligns all selected receptors through a shared class-level multiple-sequence alignment and produces a residue-level table that, for each reference position, reports the corresponding residue number and amino acid in every receptor, together with ortholog-based conservation percentages and conserved amino acids; structural region and GPCRdb generic numbering^79^ are provided for the reference receptor (Fig. 3). The full table can be downloaded as a TSV file for downstream analysis.

**Figure 3.**
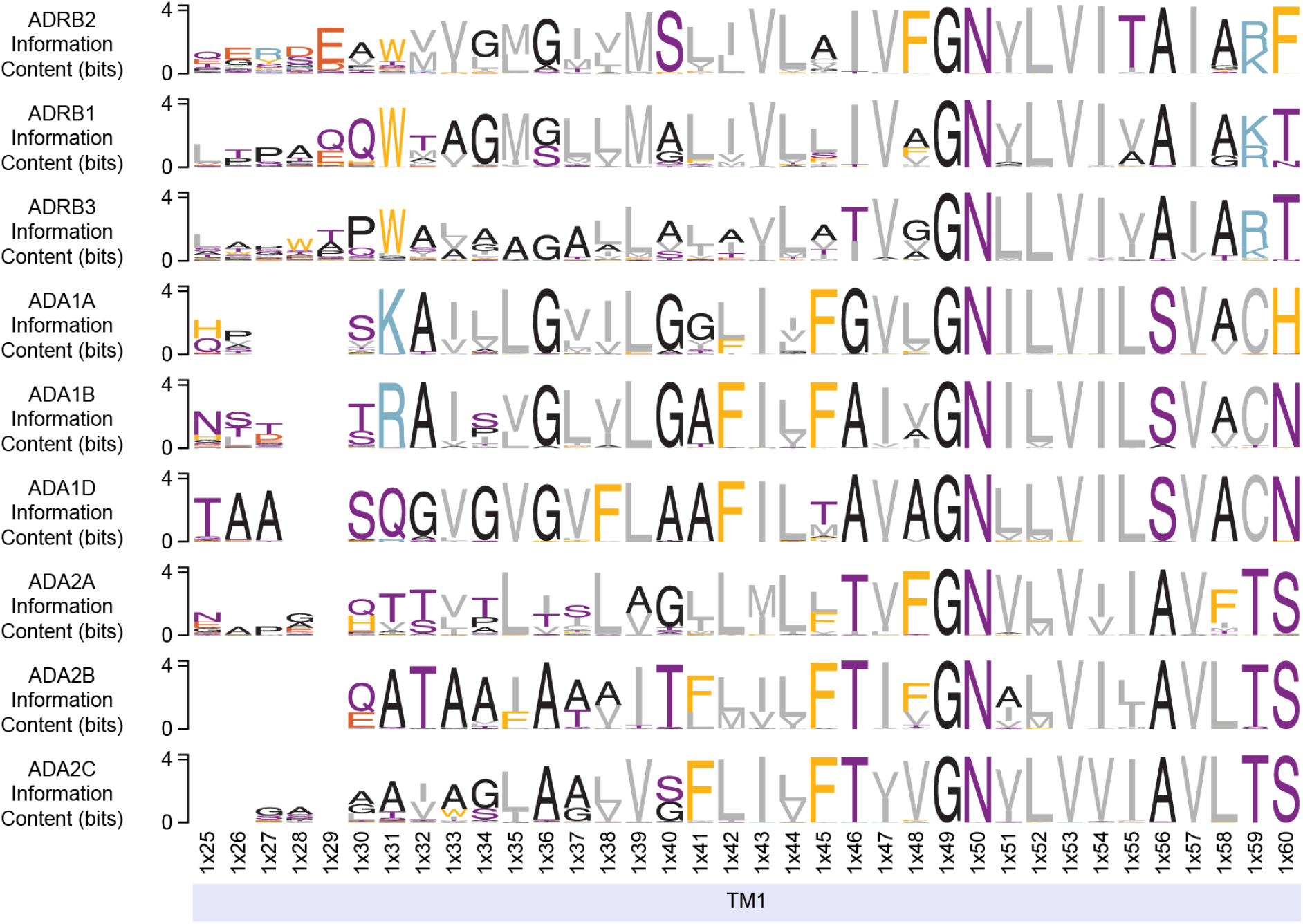
Multi-receptor comparison tool helps compare conservation patterns across multiple receptor lineages. A comparative multi-receptor sequence logo is generated by choosing Beta-2 adrenoreceptor (ADRB2) as a reference. This limits the plot to residues that can be mapped to ADRB2. For TM1, conservation patterns were compared across all adrenoceptors (ADRB1/2/3, ADA1A/B/D and ADA2A/B/C). Below the logos GPCRdb numbers^9^ for ADRB2 and structural region information is displayed.

In addition to the table, the tool generates an interactive aligned sequence logo visualization for the GPCR lineages included for comparison. For each receptor, a separate logo is built from its ortholog alignment and aligned by the reference receptor’s residue positions, allowing users to visually compare residue preferences and information content across several paralogs at once. Unlike the pairwise differential conservation tool, this interface does not automatically categorize sites as “common” or “specifically conserved”, leaving the interpretation for the user.

#### Combine Orthologs

The combine-orthologs tool lets users merge orthologous alignments from multiple human GPCRs of the same family into a single multiple-sequence alignment that includes column where either human sequence has an amino acid. After the user specifies one or more receptor gene names, the tool extracts the corresponding human sequences from family-wide MSA of human sequences, trims columns where both human sequences contain gaps, and then re-indexes each receptor’s full ortholog alignment to those trimmed human positions. For each selected receptor, the human sequence is followed by its orthologous sequences, and the resulting combined MSA is previewed directly on the website and can be downloaded as a FASTA file for downstream analyses.

Because the output is optimized for data integration rather than graphical polish, the interface includes a simple on-page alignment view while recommending that users load the FASTA file into dedicated alignment viewers such as JalView^10^ for more advanced inspection, annotation, and scripting. By harmonizing ortholog positions across several receptors, this tool enables researchers to construct custom cross-receptor MSAs that go beyond human-only comparisons and support more sophisticated evolutionary or structural analyses.

## Discussion

The GPCR Evolution Database was developed to make evolutionary analysis of GPCRs more accessible, interpretable, and useful for generating or supporting hypotheses about GPCR function. Although GPCRs are among the most extensively studied membrane protein families, research has often focused on a relatively small set of well-characterized receptors, leaving a large portion of the superfamily comparatively underexplored^11^. Our database equips researchers with high-quality evolutionary information that can help connect insights from well-studied receptors to less explored receptor groups. Furthermore, it allows existing knowledge to be reassessed from an evolutionary perspective, further enhancing our understanding of even well-studied receptor subfamilies.

Overall, the GPCR Evolution Database is an open-access platform that provides tools for exploring how evolutionary processes shape GPCR function. By enabling researchers to evaluate sequence conservation patterns within orthologs and across paralogs, the database helps identify shared ancestral features and lineage-specific residues that contribute to the functional landscape of each GPCR. In this way, it helps translate the information encoded in protein sequences over millions of years of evolution into a practical resource for improving receptor characterization.

## Methods

### Source Data

The underlying datasets used to construct the GPCR Evolution Database, including orthologous sequence sets, multiple-sequence alignments, and phylogenetic trees were retrieved from our comprehensive evolutionary analysis covering human GPCR families^8^. These datasets were subsequently processed and integrated into the database to support receptor-level visualization, residue-conservation analysis, and comparative analyses across paralogous GPCR subtypes. Generic residue numbers^7^, structural region assignments, and snake plot topologies were obtained from GPCRdb^6^.

### Calculation of Conservation Scores

Per-residue conservation was calculated from each receptor’s ortholog multiple sequence alignment using a gap-aware, similarity-based measure. For every alignment column we first determined the gap percentage. When gaps were the most frequent character at a position, that position was assigned a conservation score of 0%. Otherwise, we identified the most frequent amino acid at the position and additionally counted every other amino acid scoring greater than 1 against it in the BLOSUM80 substitution matrix, so that chemically similar residues contribute to the score. The combined count of the most frequent amino acid and its similar substitutions was then divided by the number of non-gap sequences in that column to calculate the conservation percentage. The same BLOSUM80 similarity criterion is used by the differential conservation tool to decide whether two receptors share a common residue, a position that is conserved in both paralogous lineages with a similar amino acid. Conservation scores are calculated and reported only at the alignment columns where the human receptor contains an amino acid.

## Data and Code Availability

The data and source code underlying the GPCR Evolution Database are publicly available at https://github.com/CompGenomeLab/GPCRevolution.

